# Investigating altered limbic reward system in Huntington’s Disease: Implications for apathy

**DOI:** 10.1101/2024.02.13.580195

**Authors:** Vasiliki Bikou, Inigo de Vicente, Audrey E. De Paepe, Clara Garcia-Gorro, Nadia Rodriguez-Dechicha, Irene Vaquer, Matil Calopa, Ruth de Diego-Balaguer, Estela Camara

**Affiliations:** Cognition and Brain Plasticity Unit, IDIBELL, L’Hospitalet de Llobregat (Barcelona), Spain; IDIBELL, Barcelona (Hospitalet de Llobregat), Spain; Hestia Alliance, L’Hospitalet de Llobregat (Barcelona), Spain; Movement Disorders Unit, Neurol. Service, Hosp. Universitari de Bellvitge, L’Hospitalet de Llobregat (Barcelona), Spain; Cognition and Brain Plasticity Unit, IDIBELL, L’Hospitalet de Llobregat (Barcelona), Spain / Develop. and Educ. Psychology, Barcelona Univ., Barcelona, Spain /ICREA (Catalan Inst. for Res. and Advanced Studies), Barcelona, Spain

## Abstract

Huntington’s disease (HD) is an inherited neurodegenerative condition characterized by motor, cognitive, and behavioral impairments. Apathy, marked by reduced motivation and goal-directed behavior, is the most prevalent psychiatric symptom in HD, impacting both patients and caregivers. Traditionally linked to executive dysfunction within the dorsolateral prefrontal cortex (dlPFC)-dorsal striatum loop in HD, the role of limbic regions in the basal ganglia and their impact on reward processing deficits remains poorly understood. This study sought to dissociate the functional correlates that are altered in reward valuation and may underlie apathy in HD. We aimed to tease apart whether apathy is associated with insensitivity to processing rewards, hypersensitivity to losses, or both, leading to the observed lack of motivation in apathetic individuals. Thirty-nine HD gene-expansion carriers (HDGEC) and 26 non-apathetic control participants underwent functional magnetic resonance imaging during a gambling task. The goal was to identify disrupted reward-related regions in HDGEC and their association with apathetic symptoms. Whole-brain analysis of gains and losses separately showed a significant reduction in activity, when HD group was compered to controls, within the left ventral striatum (VS), including the nucleus accumbens. The effect observed in the VS remained when clinically apathetic HDGEC were compared with controls. Conversely, non-apathetic HDGEC did not show any significant differences from controls. Interestingly, these group differences appeared exclusively during the processing of the reward. Additionally, higher levels of apathy were associated with especially decreased activity related with the processing of gains in this region. Our findings highlight the vulnerability of the left VS in HD and its association with the altered processing of gains, particularly in apathetic individuals, while preserving the valuation of losses. This suggests that reward insensitivity associated with VS dysfunction may be an important component of apathy in HD. Understanding the underlying mechanisms of reward processing and apathy in HD may help elucidate the implications of limbic regions as opposed to frontal executive dysfunction in apathy.

## 1. Introduction

Huntington’s disease (HD) is a devastating, fully penetrant hereditary neurodegenerative condition that can manifests with a range of motor, cognitive, and behavioral deficits.^1^ Neuropsychiatric abnormalities often precede motor symptoms,^2–4^ significantly impacting the quality of life for both HD gene-expansion carriers (HDGEC) and their caregivers.^5,6^ Apathy stands out as the most commonly reported behavioral deficit, with prevalence rates ranging from 46% to 76% across both pre-motor and motor manifest stages. ^7–10^

Characterized by a decline in motivation and goal-directed behavior, apathy can manifest as a dysfunction within various systems that collectively orchestrate motivated behavior.^11^ Disruptions in executive functions, including working memory, cognitive flexibility, inhibition, and planning, may result in an incapacity to effectively engage in actions towards a goal.^12^ Simultaneously, impairment in the reward system can compromise the ability to assess the rewarding and punishing potentials of actions, thereby diminishing the drive to achieving a goal.^13,14^ Apathy may emerge from the impairment of any of these processes or from the inability to integrate these complex functions.^15–17^

In the context of HD, apathy has conventionally been attributed to executive dysfunction, specifically linked with the well-documented neurodegeneration within the dorsolateral prefrontal cortex (dlPFC)-dorsal striatum loop. ^11,18^ However, there is a limited understanding of the involvement of limbic regions within the basal ganglia and their impact on reward processing deficits, which ultimately culminate in apathetic behavior. Despite this critical knowledge gap, a few studies have explored the functional correlates of apathy during the processing of rewards and punishments. Existing findings reveal divergence and contradictory trends, underscoring the pressing need for comprehensive research in this domain.^17^

Recent studies have conceptualized apathy within the framework of cost-benefit decision-making, investigating the integration of rewards and effort costs to drive goal-directed behaviour.^14^ Within this framework, apathy is understood as a disturbance of the cognitive task of evaluating potential rewards in relation to the effort or cost required to achieve them. This involves a valuation system that computes the value of a stimulus by considering its rewarding or punishing potential and modulates behavioral responses based on their associated costs, such as time and effort. This evaluation process is underpinned by a group of regions including the ventromedial prefrontal cortex, orbitofrontal cortex, ventral striatum (VS), rostral-caudate, ventral putamen and amygdala. Impairment in this system may result in an inability to estimate the value of a stimuli, hindering the ability to discriminate between favorable and unfavorable outcomes effectively. In this regard, apathetic behavior may arise from either hyposensitivity to rewards (where the effort doesn’t seem worthwhile) or hypersensitivity to punishments (where potential problems outweigh the benefits).

Existing literature does not provide a definitively conclusion regarding whether reward or punishment sensitivity is more impacted in apathy. For instance, apathetic HDGEC exhibit diminished responsiveness to positive stimuli, as indicated by challenges in recognizing socially rewarding cues compared to non-apathetic counterparts.^19^ Similar patterns are observed in related neurological conditions. Reduction of the typical pupillary response, indicative of physiological reward sensitivity, has been evidenced in clinically apathetic patient groups with Parkinson’s disease (PD) and cerebral autosomal dominant arteriopathy with subcortical infarcts and leucoencephalopathy (CADASIL) after monetary reward.^20,21^ Additionally, neuroimaging studies in PD have demonstrated compromised reward processing, as evidenced by diminished amplitude differences in feedback-related negativity potentials.^13^ Complementing these observations, studies have linked diminished activation in the orbitofrontal cortex and ventral striatum (VS) to amotivation in response to reward-related information.^22,23^ Conversely, apathetic behavior may be more closely associated with insensitivity towards punishments and negative emotional stimuli. For instance, research employing various reward-related tasks in HD patients reveals a significant correlation between apathetic symptoms and reduced responsiveness to negative outcomes or losses.^24^ This perspective aligns with prior literature emphasizing the prominent role of altered negative emotion processing in HD manifestations.^25^

In this context, we raise the critical question of how reward sensitivity is affected in HD and its putative relationship with apathy. Specifically, we aim to evaluate the reward processing system in HDGEC and investigate whether apathetic behaviour modulates brain responses associated with gains and losses. We hypothesize that apathy symptoms may be associated with insensitivity to processing rewards, hypersensitivity to losses, or both. Our exploration also extends to elucidating the role of alterations in the ventral regions of the reward network in both reward sensitivity and apathetic features in individuals with HD. To achieve this goal, we employed an fMRI reward-related task designed to localize the ventral frontostriatal network involved in the processing of the value of reward valence, and to investigate its interplay with apathy.

## 2. Materials and Methods

### 2.1. Ethics

This study was approved by the ethics committee of Bellvitge Hospital in accordance with the Helsinki Declaration of 1975 and written consent was obtained from all HDGEC.

### 2.2. Participants

Participants’ demographics and clinical data are detailed in Table 1. Thirty-nine HDGEC (44 ±3.10 CAG repeats) and twenty-six healthy, non-apathetic, control participants who were matched for age (t (55) =-0.13, P=0.98), sex (X^2^ (1, N= 65) = 0.60, P = 0.44), and years of education (t (55) = 0.66, P= 0.50) were recruited for this study. No participants reported previous history of traumatic brain injury or neurological disorder other than HD.

**Table 1:**
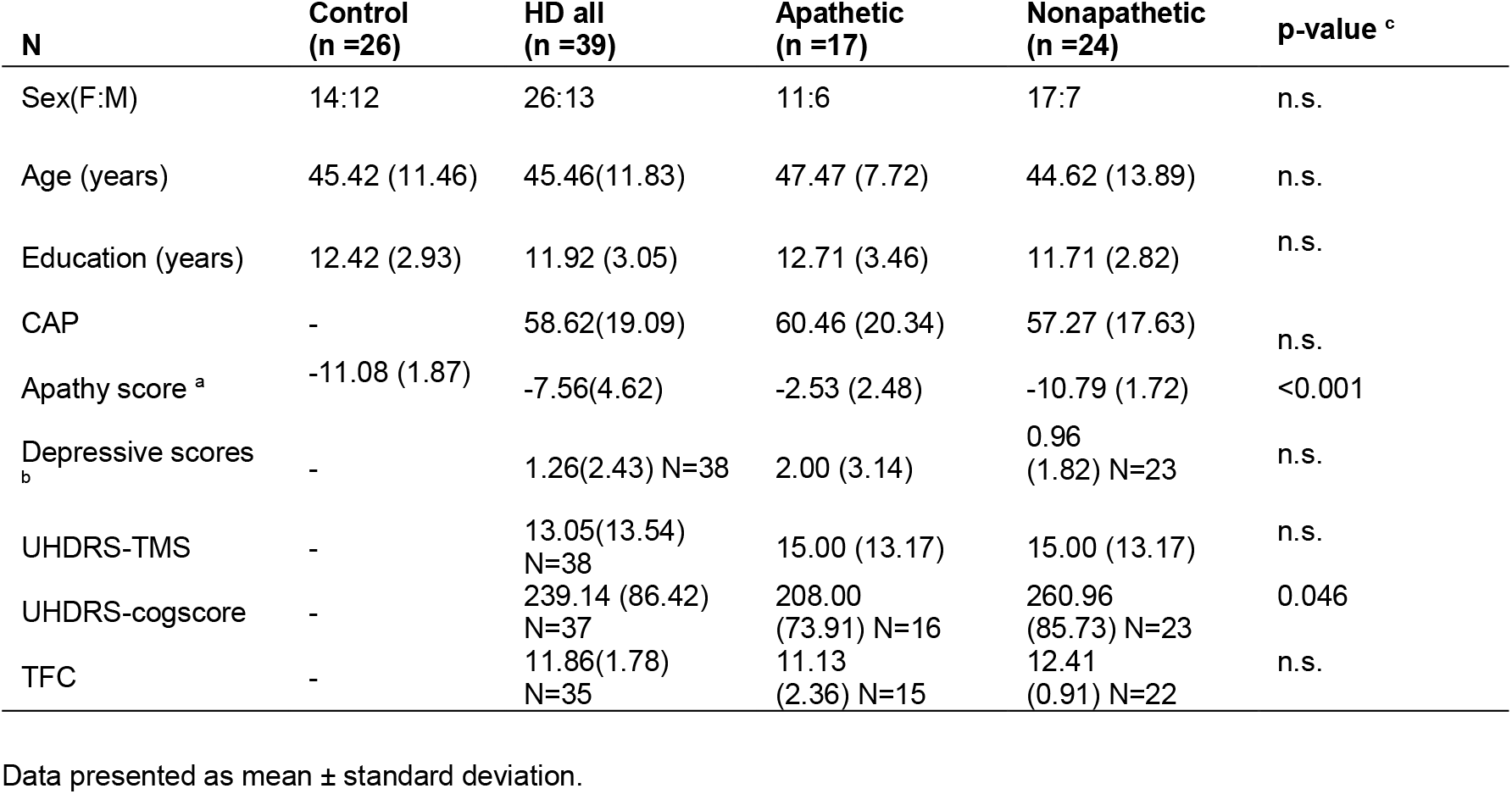
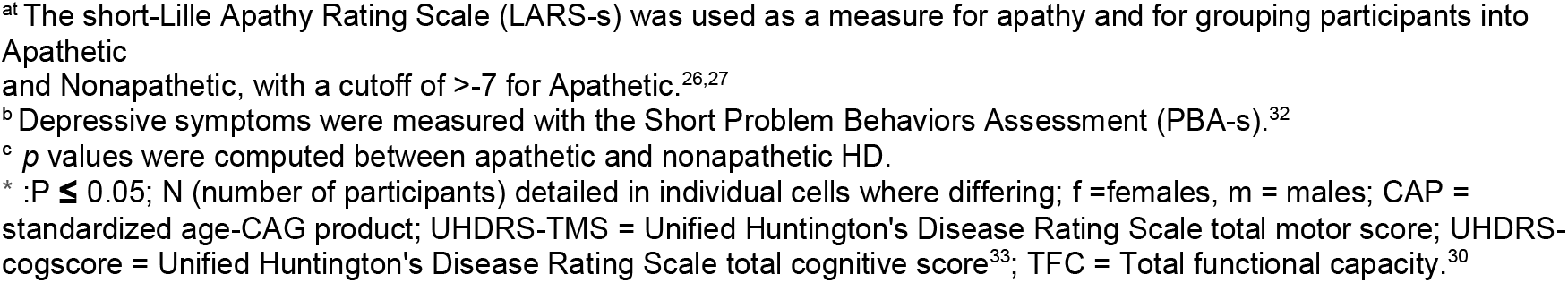
Sociodemographic and clinical characteristics of study participants.

| N | Control<br>(n =26) | HD all<br>(n =39) | Apathetic<br>(n =17) | Nonapathetic<br>(n =24) | p-value <sup>c</sup> |
| --- | --- | --- | --- | --- | --- |
| Sex(F:M) | 14:12 | 26:13 | 11:6 | 17:7 | n.s. |
| Age (years) | 45.42 (11.46) | 45.46(11.83) | 47.47 (7.72) | 44.62 (13.89) | n.s. |
| Education (years) | 12.42 (2.93) | 11.92 (3.05) | 12.71 (3.46) | 11.71 (2.82) | n.s. |
| CAP | - | 58.62(19.09) | 60.46 (20.34) | 57.27 (17.63) | n.s. |
| Apathy score <sup>a</sup> | -11.08 (1.87) | -7.56(4.62) | -2.53 (2.48) | -10.79 (1.72) | <0.001 |
| Depressive scores <sup>b</sup> | - | 1.26(2.43) N=38 | 2.00 (3.14) | 0.96<br>(1.82) N=23 | n.s. |
| UHDRS-TMS | - | 13.05(13.54)<br>N=38 | 15.00 (13.17) | 15.00 (13.17) | n.s. |
| UHDRS-cogscore | - | 239.14 (86.42)<br>N=37 | 208.00<br>(73.91) N=16 | 260.96<br>(85.73) N=23 | 0.046 |
| TFC | - | 11.86(1.78)<br>N=35 | 11.13<br>(2.36) N=15 | 12.41<br>(0.91) N=22 | n.s. |

Data presented as mean ± standard deviation.
<sup>at</sup> The short-Lille Apathy Rating Scale (LARS-s) was used as a measure for apathy and for grouping participants into Apathetic
and Nonapathetic, with a cutoff of >-7 for Apathetic.<sup>26,27</sup>
<sup>b</sup> Depressive symptoms were measured with the Short Problem Behaviors Assessment (PBA-s).<sup>32</sup>
<sup>c</sup> *p* values were computed between apathetic and nonapathetic HD.
\* : $P \leq 0.05$ ; N (number of participants) detailed in individual cells where differing; f =females, m = males; CAP = standardized age-CAG product; UHDRS-TMS = Unified Huntington's Disease Rating Scale total motor score; UHDRS-cogscore = Unified Huntington's Disease Rating Scale total cognitive score<sup>33</sup>; TFC = Total functional capacity.<sup>30</sup>

**Table 2:**
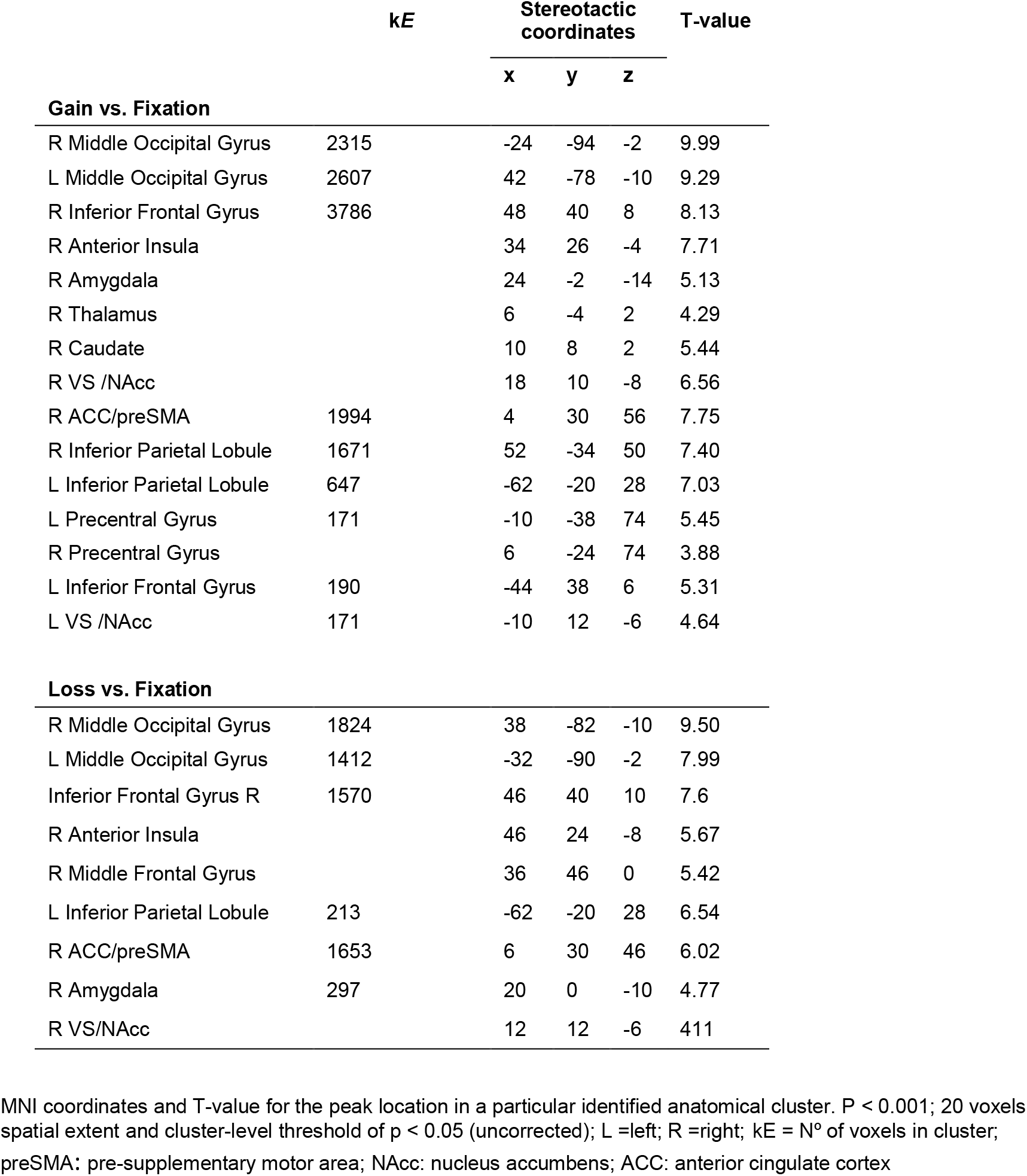
Whole sample brain activity analysis in gains and losses.

### 2.3. Clinical evaluation

The primary outcome used for this study is the Lille Apathy Rating Scale, short-form (LARS-s), an interview-based instrument that measures global apathy with scores ranging from -15 to +15, and higher scores representing a higher degree of apathy.^26,27^ Clinically relevant global apathetic syndrome is defined as a score greater than −7. ^27^ Both healthy controls and HD individuals were assessed for apathy using LARS-s. This scale is not only applicable to clinical populations but has also been effectively utilized in studies involving healthy individuals.^28,29^ To control for the effect of disease progression, the standardized CAG-Age Product (CAP) score was computed, using the form CAP = 100 × age × (CAG – 35.5) / 627.^30^ Additionally the Short Problem Behavior Assessment (PBA-s) score for depressed mood was used as a control covariate in the apathy analysis.^5,31,32^

To further characterize the HD sample, the Unified Huntington’s Disease Rating Scale for motor (UHDRS-TMS) and cognitive function (UHDRScogscore) and the total functional capacity (TFC) were also assessed and reported .^33^ All clinical assessments were carried out by neurologists or neuropsychologists specializing in movement disorders.

### 2.4. Experimental design

The experiment used a modified version of the monetary gambling task originally designed by Gehring and Willoughby.^34^ Participants were visually presented with two numbers (5 and 25), colored in black, on a screen with a white background. Two combinations could be presented: [5 25] and [25 5]. Using the left or right index fingers, the participants had to select one of the two numbers, with a maximum time of 2500ms to make the choice. The chosen number was shown underlined to confirm the bet. Subsequently, the chosen number appeared with a background either in red or green, indicating the outcome of the bet (Figure 1). If the background of the selected number turned green; it indicated a corresponding gain of the same amount in Euro Cents, whereas if it turned red, it led to a loss of that amount. The task described above constituted in two-thirds of the trials (standard trials), however, in order to enhance the reward response, in the remaining one-third, participants encountered unexpected rewards or losses (boost trials). In this condition, regardless of the chosen magnitude (5 or 25), participants either won or lost an amount of 125 Euro Cents, as indicated by a green or red background around the numbers.

**Figure 1:**
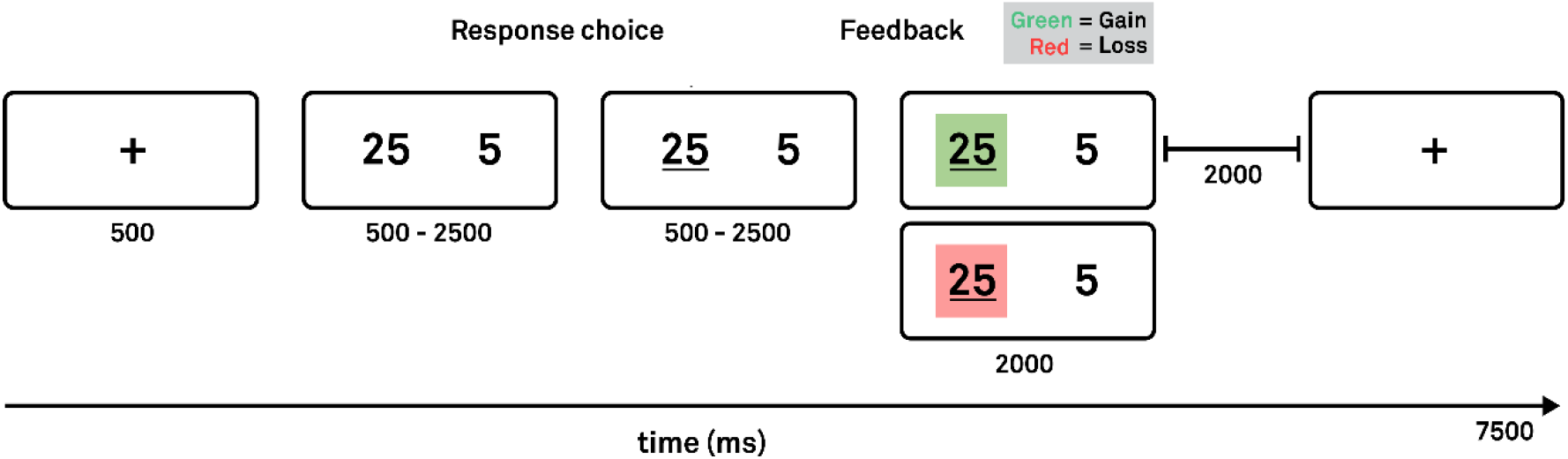
Design of the gambling task. Sequence of stimulus presentation and response events. Numbers surrounded in green represent a gain and those surrounded in red a loss.^35^

Each participant started with an initial sum of 10€ and were instructed to win to maximize their winnings. At the end of the experiment, participants received the same monetary compensation as their final score. The experiment was designed as an event-related paradigm with a total of 120 trials (40 Gain, 40 Loss and 40 Boost). Each trial was preceded by a 500ms display of a fixation cross at the screen’s center. The feedback combinations were counterbalanced and randomly presented for both standard trials ([ 5 25], [ 25 5 ], [ 5 25 ], [ 25 5 ]) and for the boost trials ([125 25 ], [ 25 125 ], [ 5 125 ], [ 125 5]). If a participant failed to press a button within 2.5 seconds during gambling, the screen displayed the text “rapido!” [fast!], and no gain or loss feedback was given for that particular trial. These trials were excluded from the analysis.

### 2.5. Neuroimaging protocol

#### 2.5.1. MRI acquisition

All MRI data was acquired on a 3T Siemens Magnetom Trio (Siemens Medical Solutions, Erlangen, Germany) at the Hospital Clinic in Barcelona while using a 32-channel head coil. Anatomical images were acquired using a high-resolution T1-weighted MPRAGE (magnetization-prepared rapid-acquisition gradient echo) sequence with the following parameters: 208 sagittal slices, TR = 1970 ms, TE = 2.34 ms, flip angle = 9°, inversion time = 1050 ms, FOV = 256 x 256 mm2, voxel size = 1 x 1 x 1 mm3. Functional images were obtained with a T2*-weighted gradient-echo imaging sequence (30 axial slices, TR = 2000 ms, TE = 30 ms, flip angle = 80º, matrix size = 64 x 64 mm2, voxel size = 4 x 4 x 4 mm3, 336 whole-brain volumes).

#### 2.5.2. fMRI preprocessing

Data preprocessing was performed using SPM12 software (https://www.fil.ion.ucl.ac.uk/spm/) and custom MATLAB scripts. After slice-timing correction to correct for the time difference between brain slice acquisitions, functional images were co-registered via an affine rigid-body transformation (i.e. translation and rotations) with the first brain volume as reference to correct for head motion. Since HD causes uncontrolled movements, motion needs special care in this clinical population. For this reason, the data were further preprocessed using ARTREPAIR (https://cibsr.stanford.edu/tools/human-brain-project/artrepairsoftware.html), which is recommended for patients prone to large motion artefacts.^37^ Following the recommended pipeline, the affine motion corrected images were smoothed with a 4mm FWHM. Gaussian kernel before repairing for motion outliers. These outliers were identified as the scans with more than 1.8% variation from the mean global signal and were repaired via interpolation to the two adjacent unaffected scans. To preserve the nature of the BOLD signal, no more than two consecutive volumes were identified as outliers were repaired. Spatial transformations were computed to co-register anatomical and functional volumes and to wrap these into the standard Montreal Neurological Institute (MNI) space. Finally, brain images were smoothed with a 7mm FWHM Gaussian kernel resulting in a total smoothing of 8mm.

### 2.6. Statistical analysis

#### 2.6.1. Sociodemographic and clinical data

Statistical analyses of group demographics and clinical variables were performed in R.^36^ Group differences of categorical variables were tested using Chi-square. Quantitative variables were described using mean and standard deviation. Normal distribution of data was inspected using Shapiro–Wilk test. Group comparisons of normally distributed demographic and clinical variables were done with student’s t test and non-normally distributed data assessed with Mann–Whitney U test (Table 1.)

#### 2.6.2. Task-related brain activation analysis

First-level analyses were based on a least-square estimation using a general linear model approach (GLM). Experimental conditions were modelled using a box-car regressor waveform and convolved using a canonical hemodynamic response function (HRF).^37^ Regressors of interest included the experimental conditions of Gain(5+25+125),Loss(5+25+125), and onsets in which participants were presented with the fixation cross were included as a baseline. To account for movement-related noise, the six rigid-body motion parameters were included as nuisance regressors in the design matrix. The data was then high-pass filtered to a maximum of 1/128 Hz and an autoregressive model (FAST) was considered in the computations to account for the temporal noise auto-correlation in the model estimates. In contrast to the default AR(1) model, the method called FAST uses a collection of exponentially decaying functions and their derivatives to fit the time series autocorrelation, which also successfully reduces false positives.^38–40^ After the estimation of the model contrast, images were generated for each participant to examine the brain activations corresponding to the experimental conditions of: Gain vs. Baseline [i.e., Gain (5 + 25 + 125) vs. Fixation] and Loss vs. Baseline [i.e., Loss (5 + 25 + 125) vs. Fixation].

First, in order to reveal the reward network engaged during the fMRI gambling task, a one-sample t-test was performed with the entire sample (i.e. Control and Huntington disease’s),. For this, the contrasts of Gain vs. Baseline and Loss vs. Baseline were used. Effects were considered significant at a whole-brain level if they exceeded a voxel-wise threshold of p < 0.001 (k > 20 voxels extent) and cluster-level threshold of p < 0.05 (uncorrected).

To identify the effect of group in each contrast one-way ANOVAs (one for Gains, one for Losses) were performed to assess the differences in activation between Controls and HDGEC.

In addition, since it is expected that the Apathetic and Nonapathetic HD subgroups differ distinctly with respect to Controls, each factor was entered separately, and then used the contrast of Controls vs. (Apathetic + Nonapathetic) to assess for the global alterations due to the disease. Results are reported at *P* < 0.001 with k > 20 of cluster extent, and cluster-level threshold of *P* < 0.05 (uncorrected).

The regions that were found to be significantly affected in HDGC were further analyzed using a ROI based ANOVA to evaluate the effect between HDGEC subgroups (Controls vs. Apathetic vs. Non-Apathetic).

To further explore the functional correlates of apathetic behavior during the processing of rewards/punishments, a multiple regression analyses was performed between apathy and fMRI activity. A whole-brain level analysis was applied for the contrast of Gain vs. baseline and of Loss vs. baseline, controlling for CAP. Correlations were considered significant if they exceeded a voxel-wise threshold of p < 0.001 (k > 20 voxels cluster extent) and cluster-level threshold of p < 0.05 (uncorrected).

#### 2.6.3. Post-hoc Analysis

Post hoc analysis at a ROI-level was conducted using the R statistical environment.^38^ The investigation into group differences involving the Apathetic, Non-Apathetic, and Control groups was carried out using one-way ANOVA tests. Subsequently, a post hoc Tukey analysis was applied to identify specific group differences. Differences were considered statistically significant when P-adj ≤ 0.05, the Tukey-Kramer method was employed for multiple testing correction to control the false discovery rate. Spearman correlations were performed to assess the relationship between apathy severity scores and effect sizes. To examine apathy as an independent psychiatric syndrome from depression and account for the disease progression, the PBA-s ‘depression’ component, CAP and age were utilized as covariates of no interest.^41,42^

## 3. Results

### 3.1. Demographic and clinical data

Demographics and clinical characteristics of participants are listed in Table 1. As expected, the apathetic group demonstrated significantly higher apathy scores than the nonapathetic group (Mann–Whitney U= 408, n1= 17, n2= 24, *P*<0.0001). Additionally, when apathetic and nonapathetic HD participants were pooled together, the group maintained significant differences compared to the control group in the global LARS-s group (Mann–Whitney U= 822, n1= 41, n2= 35, p=0.015). Comparisons between apathetic and nonapathetic HD individuals in the rest of the clinical measures revealed no group differences other than UHDRS cognitive scores, a metric of global cognitive functioning, with the apathetic group having lower total scores (t (35) = -2.0, *P*= 0.046). Depressive symptoms were non-significantly lower in nonapathetic compared to apathetic HDGEC. Notice that there were no significant differences in motor disease severity or CAP between groups.

### 3.2. Neural correlates of Reward Valuation

To identify the network supporting reward valuation we compared whole-brain activity when participants (*i*.*e*., both control and HD groups together) obtained a monetary outcome (*i*.*e*., Gains and Losses) against Fixation. For both monetary gain or loss conditions, this analysis revealed overlapping fronto-subcortical-limbic-parietal networks. As expected, significant enhanced hemodynamic activity were observed in the ventral striatum (NAcc) bilaterally (with the activity extending to the caudate, the amygdala and the insular context), the prefrontal cortex (the anterior cingulate cortex, the inferior and middle frontal cortex), the inferior parietal cortex, and the middle occipital gyrus (Fig. 2).

**Figure 2:**
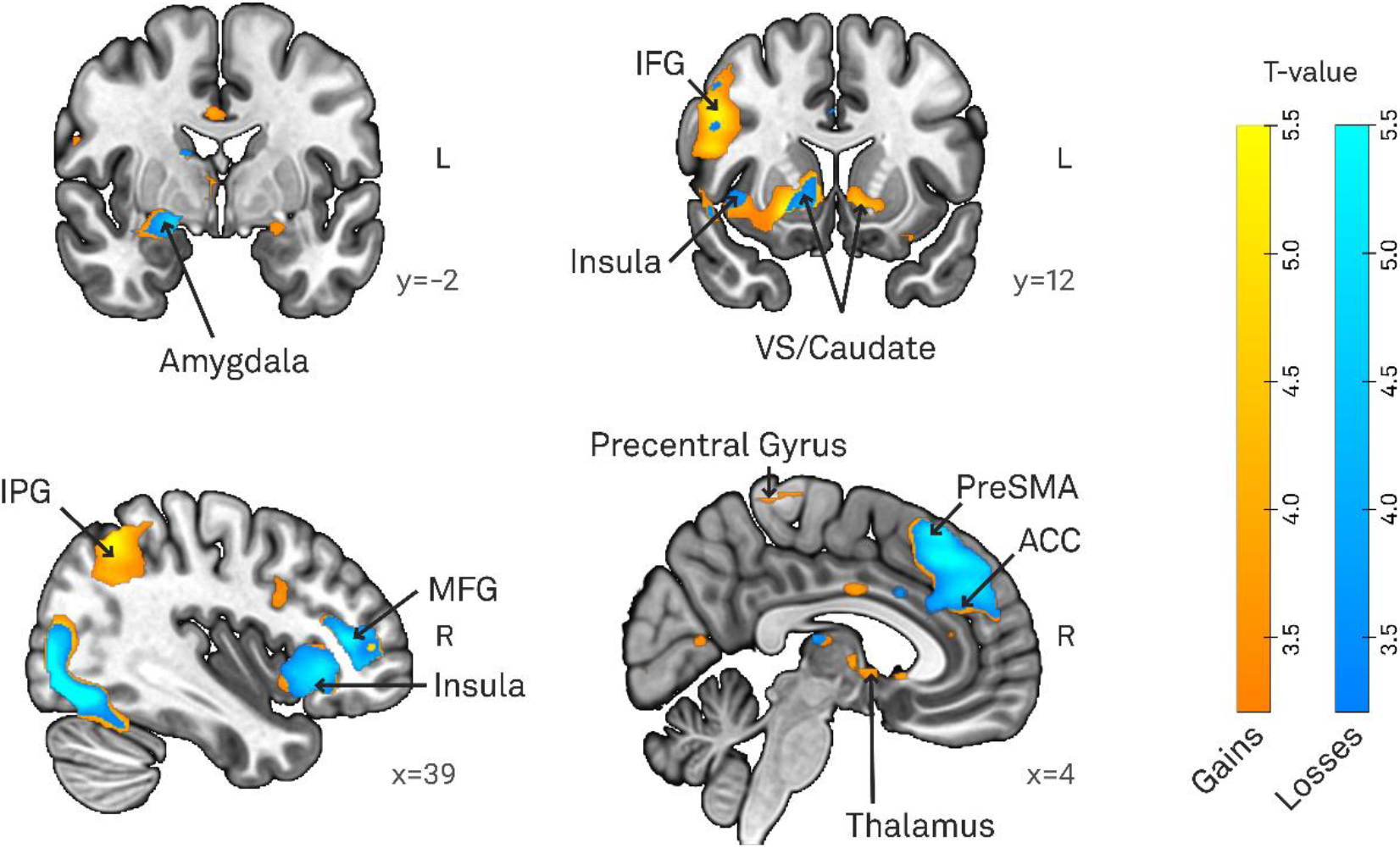
Common activations in both Control and HDGEC groups for the gains and losses. Coronal and sagittal views of whole-brain univariate functional analysis (HD and Controls). Cold and warm colours refer to activation in loss vs. fixation contrast and gain vs. fixation contrast, respectively. Statistical maps are superimposed onto the MNI structural template. IFG Inferior Frontal Gyrus; IPG: Inferior Parietal Lobule; MFG: Middle Frontal Gyrus; ACC: anterior cingulate cortex; preSMA: pre-supplementary motor area

### 3.3. Group differences in Gain and Loss Valuation

The voxel-wise group comparison revealed reduced levels of activity in the limbic regions of the reward valuation network when comparing the HDGC group with healthy controls. Specifically, during the processing of gains, HDGC presented significantly lower levels of activity in the left ventral striatum (including NAcc) and ventral parts of caudate and putamen (peak activity, MNI coordinates, x, y, z, left hemisphere, −12, 16, −2, *T* = 3.2, *P* < 0.001; *P*-value at cluster level uncorrected; Fig. 3A). When examining the response to monetary loss trials, significant differences were found only after lowering the threshold to *p* < 0.005 voxel-level (still with *p* < 0.05 cluster level). Increased activity levels were observed in the HD group compared to the controls within the right dlPFC (peak activity, MNI coordinates, x, y, z, right hemisphere, 42, 24, 42, *T*=2.6, *P* < 0.005; at cluster level uncorrected; Fig. 3D).

**Figure 3:**
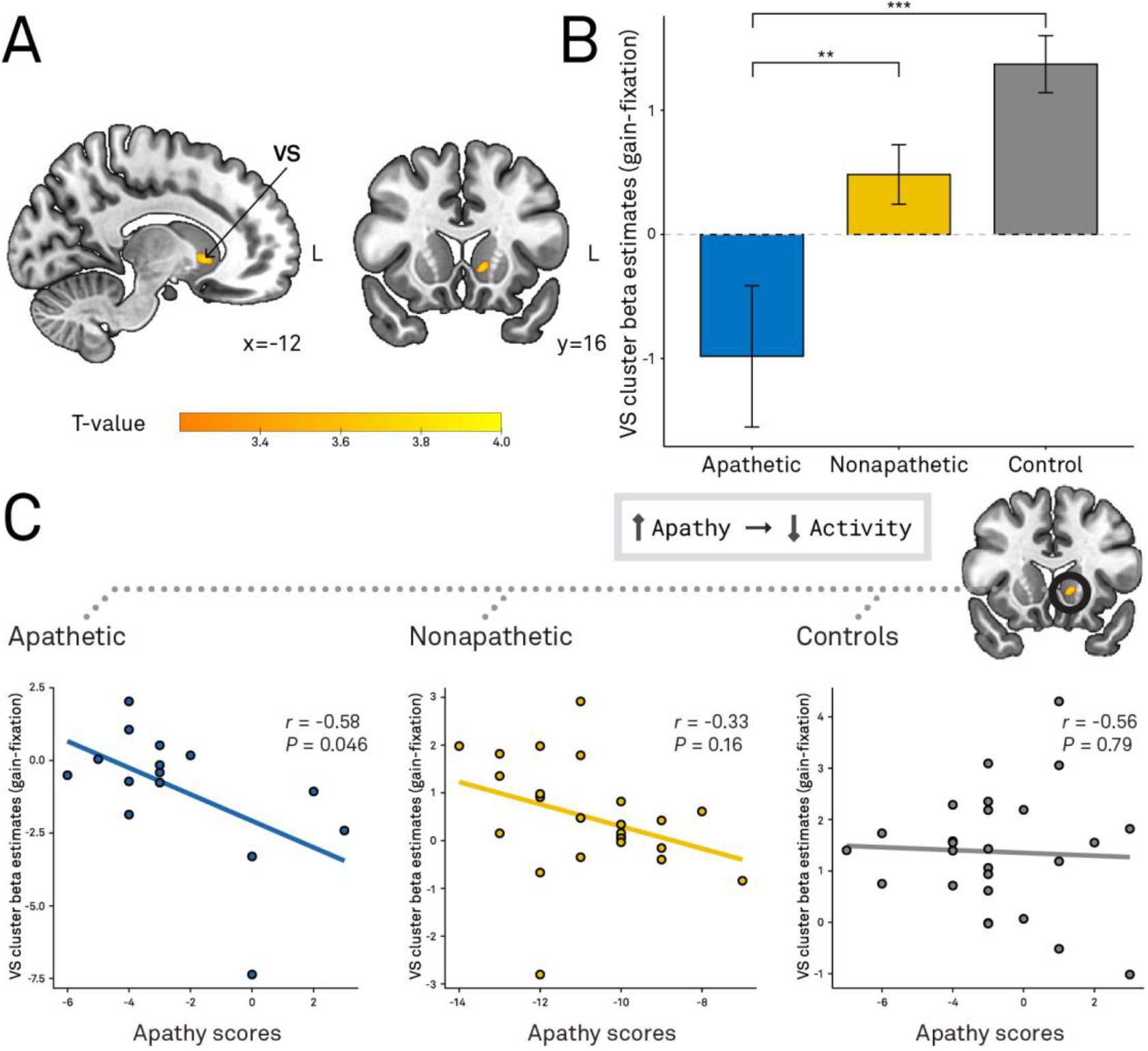
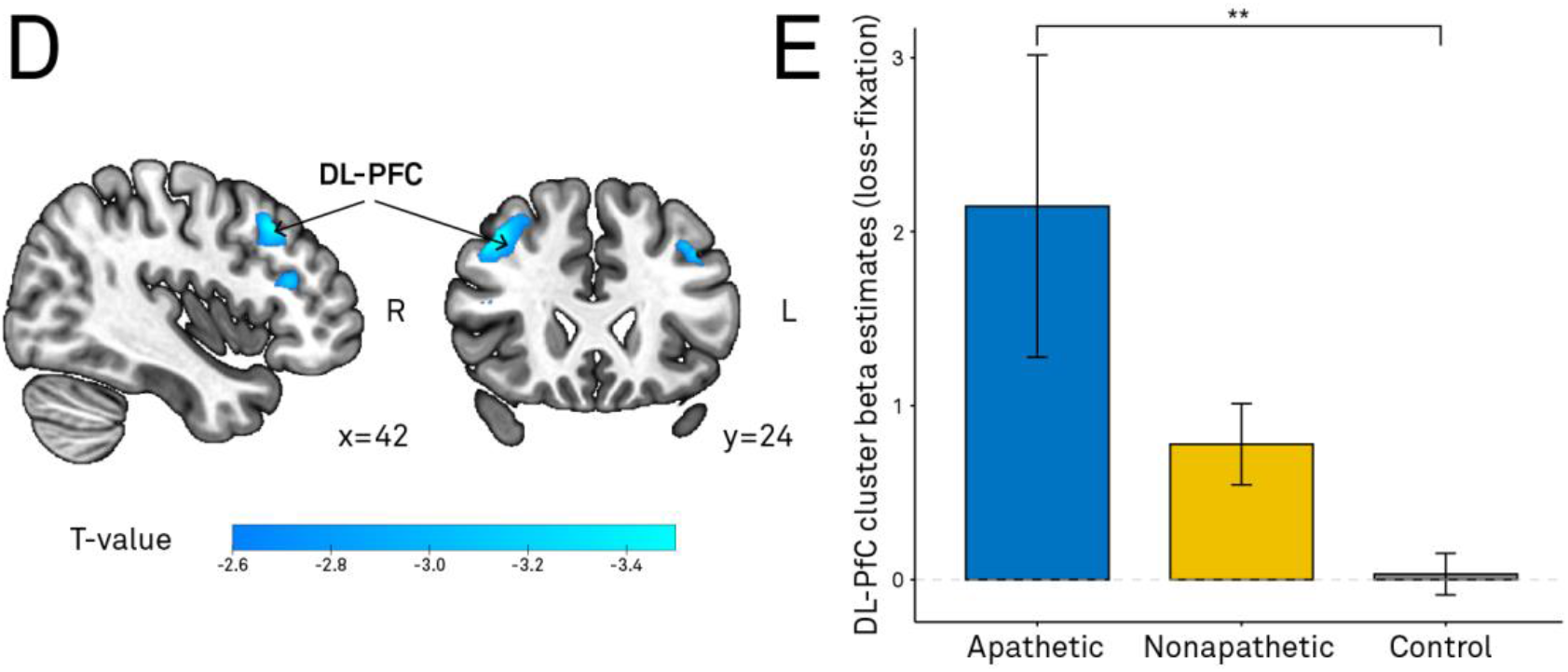
Differences in brain activity between Controls and HDGEC. (A) Significantly higher brain activity in dlPFC in the HD group compared to the controls. (B) From the significant contrast shown in C, raw data-points per individual have been extracted from mask of this cluster, to demonstrate HD subgroup differences. The barplot represent the average activity found for each group: Apathetic (left), nonapathetic (middle), controls (right). Error bars indicate the standard error of the mean. (C) Significantly higher brain activity in the nucleus accumbens, caudate nucleus and putamen in the control group compared to the HDGEC. (D) Barplot was created similarly to B. (E) Relationship between apathy levels and brain activity in the ROI in the three groups of interest. Bivariate plot displaying the significant association between global apathy and beta values extracted from VS ROI. Apathy is measured with the short-form Lille Apathy Rating Scale (LARS-s), ranging from −15 to +15, where larger and more positive scores indicate more severe apathy. Correlation coefficients (r) and raw P-values (P) for each group are shown in upper right. Linear regression line is fit for each scatterplot to aid interpretation. NAcc: Nucleus accumbens; * :P < 0.05; ** P ≤ 0.01; ***: P<0.001; **** :P ≤ 0.0001 after controlling forI multiple comparisons.

### 3.4. Relationship between altered reward processing and apathy

Based on the significant effects observed in the left ventral striatum and the right DLPFC, post-hoc analyses at an ROI-level were performed to analyze the effect between HDGEC subgroups (Controls vs. Apathetic vs. Nonapathetic). As expected from the group activation maps, we found significant group differences in in ventral striatum during gain trials (one-way ANOVA; *F* (2,62) =12.16, *P*<0.001) and in dlPFC during loss trials (one-way ANOVA; *F*(2,62)=6.7, *P*=0.002). This difference was driven by lower levels of activation in the *apathetic* group in ventral striatum (post hoc Tukey: apathetic versus controls: mean difference= 2.4, *P*< 0.001; non-apathetic versus apathetic: mean difference= 1.5, *P*= 0.01; controls versus non-apathetic: mean difference= -0.9, *P*=0.91; Fig. 3B) and higher levels of activity in apathetic individuals in dlPFC (post hoc Tukey: apathetic versus controls: mean difference=-2.1, *P*= 0.002 non-apathetic versus apathetic: mean difference= -1.4, *P*= 0.059; controls versus non-apathetic: mean difference= 0.7, *P*=0.308; Fig. 3E).

The relationship between the beta values of the left ventral striatum and apathy measures was examined using Spearman partial correlations, while controlling for depression, CAP, and age. Among all HD individuals (apathetic and nonapathetic combined), a significant negative correlation was found between beta values and scores from the LARS-s (*r* = -0.51, *P* =0.002). Isolating the apathetic HDGEC resulted in a similar significant result (*r*=-0.58, *P*=0.046). Conversely, no statistically significant correlations were found in the control or the group of nonapathetic HD individuals. (Fig. 3C). No relationship was found between levels of brain activity in dlPFC and apathy scores in any of the groups.

### 3.5 Apathy correlates in Gain and Loss Valuation

To explore individual differences in apathy, after controlling for depression and CAP, and their potential impact on the reward valuation system, we employed a whole-brain approach. Our objective was to investigate brain regions not exhibiting functional deficits but possibly involved in the manifestation of the spectrum of apathy. Our focus was on the HDGEC group, given the variability in apathy scores within this cohort. Higher levels of apathy were associated with decreased brain activity, during monetary gain trials in a fronto-striatal-limbic network, including the left caudate, the bilateral insular cortices, the left superior and middle temporal gyri(Fig. 4A; Table 3). No such correlations were observed during monetary losses.

**Table 3:**
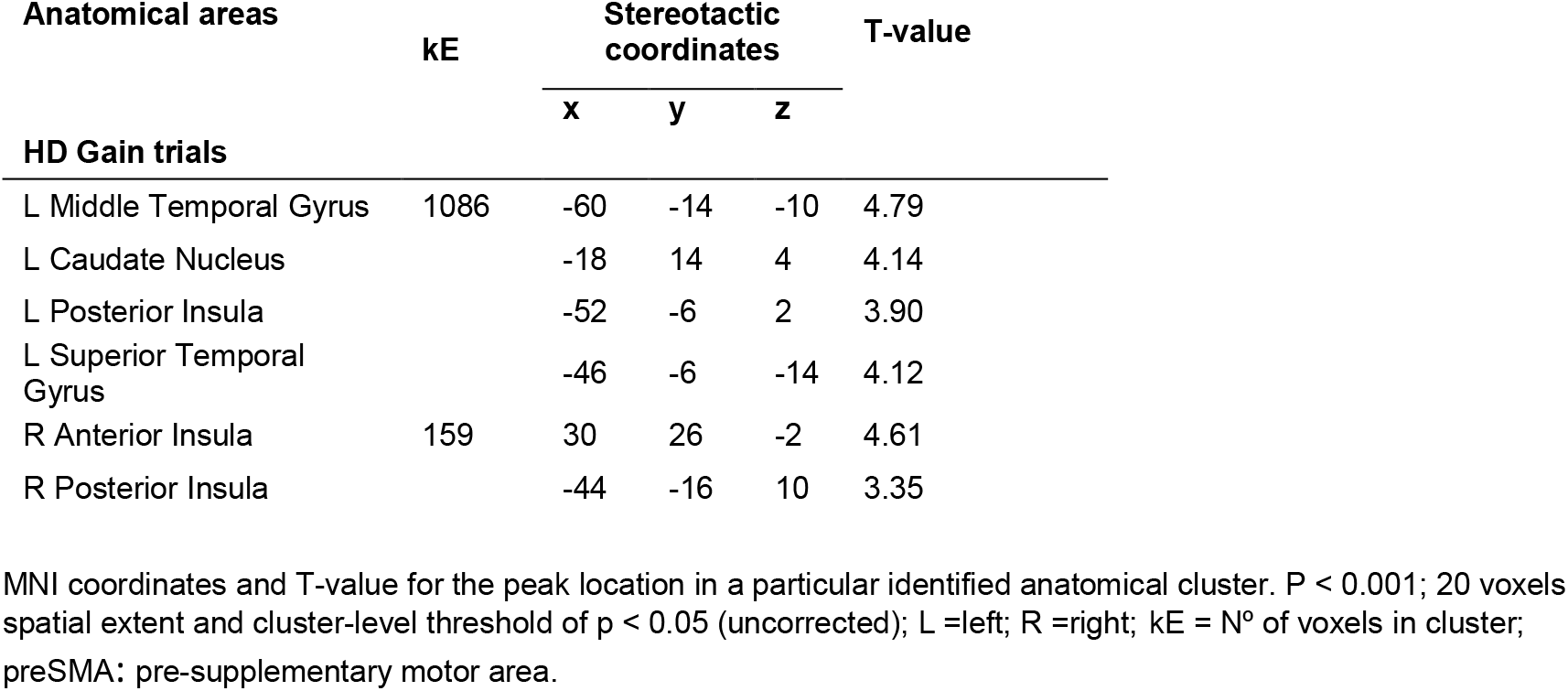
Correlations between apathy and brain activity.

| Anatomical areas | kE | Stereotactic coordinates |  |  | T-value |
| --- | --- | --- | --- | --- | --- |
|  |  | x | y | z |  |
| HD Gain trials |  |  |  |  |  |
| L Middle Temporal Gyrus | 1086 | -60 | -14 | -10 | 4.79 |
| L Caudate Nucleus |  | -18 | 14 | 4 | 4.14 |
| L Posterior Insula |  | -52 | -6 | 2 | 3.90 |
| L Superior Temporal Gyrus |  | -46 | -6 | -14 | 4.12 |
| R Anterior Insula | 159 | 30 | 26 | -2 | 4.61 |
| R Posterior Insula |  | -44 | -16 | 10 | 3.35 |

MNI coordinates and T-value for the peak location in a particular identified anatomical cluster. $P < 0.001$ ; 20 voxels spatial extent and cluster-level threshold of $p < 0.05$ (uncorrected); L =left; R =right; kE = N<sup>o</sup> of voxels in cluster; preSMA: pre-supplementary motor area.

**Figure 4.**
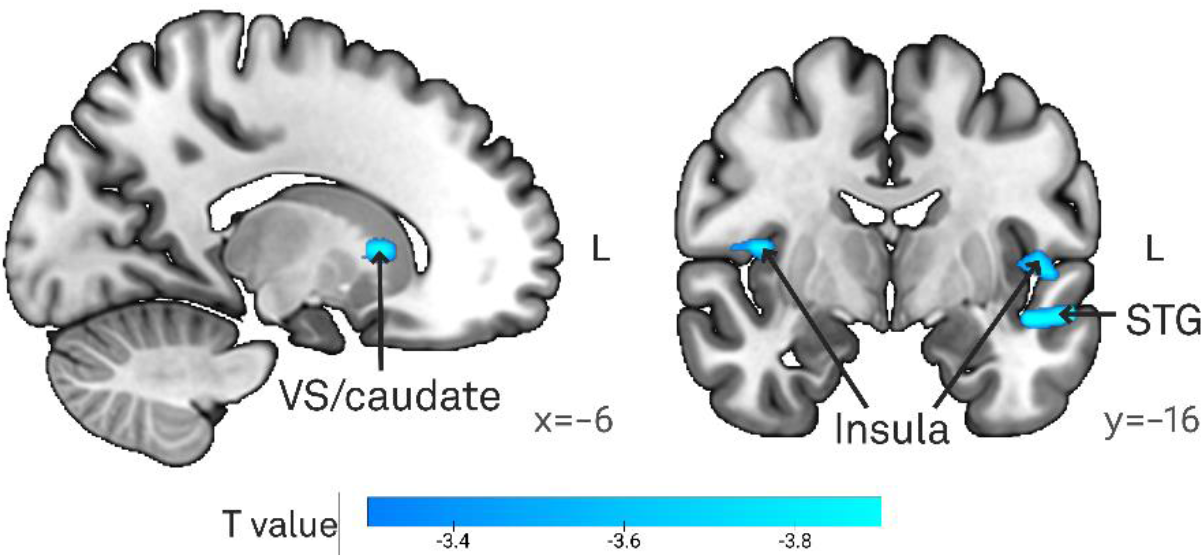
Brain regions associated with apathy scores during the possessing of bet outcome.

## 4. Discussion

This research sought to evaluate the functionality of the reward-valuation system in individuals with HD and investigate how apathetic behaviour influences brain signalling related to gains and losses in a gambling task. First, utilizing an fMRI reward-related task enabled the targeting of the limbic parts of the frontostriatal loop, which evolves key areas essential for goal-directed behaviour, emotional processing, mood regulation, and affect.^43^ Employing a whole brain analysis approach, we identified specific brain regions affected in HD participants during the evaluation of gains and losses. Our findings indicate that HD individuals exhibit diminished activity in the left VS specifically during gain outcomes. This lateralization pattern has previously been reported in HD with the left striatum being predominately affected.^16,44^ Further, upon examining signal alterations at an ROI level, we assessed activity disparities among distinct HDGEC subgroups (Controls vs Apathetic vs Nonapathetic). Results revealed that those categorized as apathetic exhibited especially reduced activity levels compared to both the nonapathetic and control groups. Interestingly, a direct and significant association emerged between the severity of apathy and VS activation, wherein diminished activity correlated with heightened apathy scores, a pattern observed within the apathetic group. This observed relationship remained consistent irrespective of disease progression or depression severity, exemplifying that apathetic behaviour selectively modulates VS response to gains.

Additionally, during loss processing, HD participants manifested heightened activation in the right dlPFC. Subsequent ROI-based analyses revealed that this distinction was driven from the apathetic group relative to the control group. Unlike the findings related to the VS activity, no direct modulatory effect of apathy was evident. Lastly, when investigating the individual differences in apathy focusing on HDGEC, we observed that heightened apathy scores corresponded with decreased brain activity during monetary gain trials in the bilateral insular cortices, the left superior and middle temporal gyri, and the left caudate nucleus. This investigation highlights brain regions that may not directly exhibit functional neuroimaging alterations in the HD, but rather contribute to apathetic manifestations.

Accumulating evidence points to the VS (including NAcc) as an important region of the network that underlies goal-directed behaviour.^14,45^ In our study, focusing on the outcome or feedback phase of the gamble (rather than the anticipation) evidenced reduced left VS signal during gains in HD apathetic individuals. Such results correlated with an increase in apathy severity. It is plausible that the activity observed during this phase signifies prediction error signal, reflecting the disparity between the expected and actual rewards, especially in situations where contingencies are uncertain.^46,47^ These signals play a crucial role in facilitating learning about which behaviors hold value for future performance, increasing the likelihood of a behaviour that leads to a better-than-expected outcome will be repeated.^48,49^ Interference with this mechanism can result in a learning bias, altering the balance between the assessment of costs or losses and rewards or gains. Over time, this bias may distort the perception of whether an action is worthwhile, ultimately leading to a decline in motivation and apathetic symptomatology. Observing this functional deficit specifically during the processing of gains may suggest an impairment in handling rewarding outcomes, potentially leading to compromised learning from rewards. Comparable findings have surfaced in studies involving Parkinson’s disease, cerebral small vessel disease, and schizophrenia, where researchers identified a connection between apathy and decreased reward sensitivity.^13,15,20,22^ In the context of HD, the literature exhibits certain inconsistencies. There is evidence that valuation and learning mechanisms are affected on both sides of the spectrum, that is, for reward and punishment.^24,44,50,51^ However, to our knowledge, there are no studies that have directly linked these outcomes to apathy and its neuronal correlates. We propose a possible link between the functional role of the VS in outcome processing to the well-known early striatal degeneration as a possible neuronal mechanism of apathy in HD.

In contrast to the observed responses of the VS to positive reinforcement, the dlPFC signaling to punitive signals was less expected. While it is not uncommon to detect dlPFC activation after punitive outcomes,^52–54^ the experimental design of this study is tailored to stimulate limbic regions associated with reward processing, rather than cortical components. Notably, the increased dlPFC activation observed in the HD group appears to be specific to punitive consequences. One possible interpretation for this activation pattern lies in the potential anatomical differentiation of outcome valence. Although debated, there is evidence of distinct neural substrates for processing rewards and punishments. Specifically, the anterior insula and dlPFC have been found to respond mainly to negative outcomes, while a network including ventromedial prefrontal cortex, VS, posterior cingulate cortex and ventrolateral orbitofrontal cortex responds mainly to positive outcomes.^47,52,54–58^ From a functional perspective, the involvement of the dlPFC in motivational processes could indicate the existence of an attentional system that regulates ongoing behavior, facilitating shifts or interruptions in response to punitive cues. Such a conceptualization aligns with the well-established cognitive control and inhibitory functions attributed to the dlPFC.^59^

In the context of our findings, the increased dlPFC activity observed within the apathetic HD group may be indicative of several underlying mechanisms. Firstly, it could be understood as a compensatory neural response, suggesting an attempt by the residual neuronal populations to offset the neurodegenerative impact, particularly given the known vulnerability of the dlPFC in early stages of the disease progression.^60^ Alternatively, this heightened activity might reflect an enhanced sensitivity to punitive stimuli, that can lead to response inhibition or avoidance behavior.^61^ However, the absence of a direct association between brain activity and apathy in this specific region raises questions about the potential contribution of this functional imbalance to the neurobiology of apathy.

The final objective of this study is to identify regions that, while not necessarily functionally impacted in HD, may be linked to apathetic symptoms. For this, we investigated the individual differences in apathy inside the HD group while controlling for depression and disease progression. As expected, we replicated the previous result of the modulation in VS and caudate nucleus. Additionally, we observed that the insula cortex and temporal lobe model the severity of apathy in HD. Prior research has also highlighted these regions in relation to apathy across various conditions.^12^ In the context of HD, Martinez-Horta *et al*. emphasized a strong correlation between apathy severity and structural impairments in the temporal cortex, suggesting that the functional deficits we identify may stem from reductions in gray matter volume.^62^ Similarly, Stanton *et al*. found diminished gray matter volume in the dorsal anterior cingulate cortex and insula among apathetic HDGEC with progressive supranuclear palsy or Alzheimer’s disease, reinforcing the implication of insular cortex.^63^ A possible explanation for these regions not being as consistently related to apathy could be their association with distinct dimensions of apathy. For instance, in Alzheimer’s disease and frontotemporal dementia, decreased gray matter intensity of insular cortex was specifically linked with affective apathy, not necessarily cognitive or behavioral.^12^ These observations illustrate a complex interaction of multiple brain changes underlying apathy in HD that evolve regions outside the basal ganglia and prefrontal cortex.

The present study is not without limitations. Firstly, our focus on HD patients in the early and intermediate stages can limit the applicability of our findings. Future research should broaden its scope to include later disease stages to better understand the progression of reward-related impairments and apathy. This is crucial, as it is possible that different mechanisms are responsible for behavioral changes as the pathology of the disease develops in time.

Additionally, prior studies have associated apathy with effortful actions,^29,64,65^ suggesting a need for investigating apathy within more natural, effort-demanding decision-making tasks. Unlike our study, which employed a straightforward binary choice paradigm with no inherent advantages or disadvantages, exploring apathy’s impact in situations requiring more complex decision-making processes could yield insightful outcomes. Nonetheless, our paradigm specifically targets limbic regions within the reward network, offering valuable insights into the relationship between alterations in the ventral part of striatum and apathy. We emphasize the involvement of regions beyond the basal ganglia in contributing to the expression of apathy highlighting the importance for future studies to explore the interactions between various systems involved in different aspects of motivation. In this pursuit, recognizing the different subtypes of apathy becomes essential, as distinct dimensions of apathy may correspond to diverse neuronal signatures.^18^

In conclusion, the present study sheds light into the neuronal mechanisms underlying apathetic behavior in HD. We found abnormal activity in VS during the processing of favorable outcomes which can reflect disrupted valuation system and result to reduced sensitivity to reward in apathetic HDGEC. Considering the importance of apathy in so many disorders, we consider fundamental to understand the cognitive mechanisms that result in this motivational impairment. A better understanding of the impact of rewards and punishments in this population, and their functional correlates, could help develop new therapeutic strategies to improve their day-to-day lives.

## Acknowledgements

The authors are grateful to the patients and their families for their participation in this project. We would also like to thank Dr. Saül Martinez-Horta, Dr. Jesús Pérez Pérez, Dr. Jaime Kulisevksy, Pilar Sanchez, Dr. Esteban Muñoz, Celia Mareca and Dr. Ruiz-Idiago for help with clinical evaluation.

